# TMS reveals a two-stage priming circuit of gesture-speech integration

**DOI:** 10.1101/2023.04.04.535555

**Authors:** Wanying Zhao

**Author notes:** Correspondence: Dr. Wanying Zhao.

## Abstract

Naturalistically, multisensory information of gesture and speech is intrinsically integrated to enable coherent comprehension. Such cross-modal semantic integration is temporally misaligned, with the onset of gesture preceding the relevant speech segment. It has been proposed that gestures prime subsequent speech. However, there are unresolved questions regarding the roles and time courses that the two sources of information play in integration. Here, in two between-subject experiments of healthy college students, we segmented the gesture-speech integration period into 40-ms time windows (TWs) based on two separately division criteria, while interrupting the activity of the integration node of the left posterior middle temporal gyrus (pMTG) and the left inferior frontal gyrus (IFG) with double-pulse transcranial magnetic stimulation (TMS). In Experiment 1, we created fixed time-advances of gesture over speech and divided the TWs from the onset of speech. In Experiment 2, we differentiated the processing stages of gesture and speech and segmented the TWs in reference to the speech lexical identification point (IP), while speech onset occurred at the gesture semantic discrimination point (DP). The results showed a TW-selective interruption of the pMTG and IFG only in Experiment 2, with the pMTG involved in TW1 (−120∼-80 ms of speech IP), TW2 (−80∼-40 ms), TW6 (80∼120 ms) and TW7 (120∼160 ms) and the IFG involved in TW3 (−40∼0 ms) and TW6. Meanwhile no significant disruption of gesture-speech integration was reported in Experiment 1. We determined that after the representation of gesture has been established, gesture-speech integration occurs such that speech is first primed in a phonological processing stage before gestures are unified with speech to form a coherent meaning. Our findings provide new insights into multisensory speech and co-speech gesture integration by tracking the causal contributions of the two sources of information.

## Introduction

In communication, messages can be expressed through multiple modalities, during which information of all kinds are treated equally regardless of the modality from which the information comes (MacSweeney et al., 2004; Hagoort and van Berkum, 2007; Weisberg et al., 2017). Moreover, naturalistically, the occurrence of information from two modalities is not neatly aligned (for a review, see Holler and Levinson, 2019), and prediction is considered to be fundamental for the processing of this temporally misaligned information (for a review, see de Lange et al., 2018)

As extralinguistic information, gestures have often been observed to accompany language. One particular type of gesture is the iconic gesture, which not only conveys relevant information but also additional information that is not present in the accompanying speech. For example, by moving the fingers in an inverted V shape while saying, “He walked across the street”, or by making an upward climbing movement with the hands when saying, “He climbs up the wall”. In the present study, when referring to gestures, we specifically meant iconic gestures. Gestures are so commonly co-occurring with speech that speakers convey information in both gesture and speech, while listeners are sensitive to information made available in both modalities (Kelly and Church, 1998; Goldin-Meadow and Sandhofer, 1999). During speech, related gestures function as a communicative device such that people have better comprehension when they are presented with speech accompanied by gestures than when they are presented with speech alone (Kelly, 2001; Valenzeno et al., 2003; Hostetter, 2011; Hostetter and Alibali, 2011). In fact, from a functional point of view, gestures can be regarded as ‘part of language’ (Kendon, 1997) or functional equivalents of lexical units in spoken language to alternate and integrate with speech.

Temporally, gestures have been supposed to occur ahead of (Morrel-Samuels and Krauss, 1992) or even terminate before (Fritz et al., 2021) their semantic affiliate. Both temporal relationships, i.e., gestures leading speech (Kelly et al., 2004; Wu and Coulson, 2005; Holle and Gunter, 2007; Ozyurek et al., 2007; Kelly et al., 2010a; Yap et al., 2011; Pine et al., 2013) and gestures occurring partly ahead of the speech segment (Kelly et al., 2010a; Drijvers et al., 2018; Zhao et al., 2018; Zhao et al., 2021; Zhao et al., 2022), have been examined in previous research. However, the question of how gestures contribute to gesture-speech integration remains unanswered, despite various methodological investigations being carried out (for a review, see Kandana Arachchige et al., 2021).

Studies have been conducted to explore the time phase for the effect of gestures on speech by manipulating the degree of synchronization between gestures and speech. Habets et al. (2011) discovered that the N400 effect, which illustrates a priming effect of gesture over speech in the semantic phase, was found when a gesture was presented either synchronously or 160 ms ahead of speech. Additionally, Obermeier et al. (2011) replaced the full gesture and speech with fragments of minimal length required for semantic identification, i.e., the discrimination point (DP) of gestures and the identification point (IP) of speech, to investigate the synchronization of gesture-speech presentation. By manipulating the temporal alignment of the homonym and the gesture, Obermeier and Gunter (2015) reported a significantly triggered N400 effect when gestures were presented from −200 ms to +120 ms in reference to the IP of speech. More recently, a transcranial magnetic stimulation (TMS) study described a neural circuit underpinning the prelexical sensory processing stage and the postlexical processing stage of gesture-speech integration by aligning speech onset with gesture DP, thus creating a semantic priming effect of gesture upon speech and segmenting eight time windows (TW) in accordance with speech IP (Zhao et al., 2021).

Based on all the above studies, we conclude that there is no consensus regarding how gestures prime speech. Some researchers have suggested that gestures prime speech in the phonological phase (Rauscher et al., 1996; Hadar and Butterworth, 1997; Krauss, 2000), some have found evidence only for the semantic phase of speech processing (Wu and Coulson, 2005; Holle and Gunter, 2007; Ozyurek et al., 2007), others have found an effect in both phases (Kelly et al., 2004), and still others have not discriminated between the two phases (Habets et al., 2011; Obermeier et al., 2011; Obermeier and Gunter, 2015).

In two experiments, the present study aimed to settle the debate by segmenting the gesture-speech integration period into scales of different referential. Experiment 1 referred to the time course of the integration, where speech onset occurred at 200 ms after the gesture stroke, and the TWs were segmented from the onset of speech (Kelly et al., 2010a; Zhao et al., 2018) (**Figure 1A**). Experiment 2 referred to the processing stage of gesture-speech information, with speech onset occurred at the DP of gestures while the TWs were divided relative to the IP of speech (Zhao et al., 2021; Zhao et al., 2022) (**Figure 1B**). Online double-pulse TMS stimulation, a method that is ideal for examining causal brain-behavioral relationships, was applied over either the left IFG or the left pMTG, brain locations believed to be responsible for gesture-speech integration (Zhao et al., 2018; Zhao et al., 2021). Previous studies have shown an inhibitory effect on cortical functions when TMS double pulses were applied with a pulse interval of 40 ms (O’Shea et al., 2004; Pitcher et al., 2008; Zhao et al., 2021). Therefore, time windows of 40 ms were split separately in the two experiments. We hypothesized that TMS stimulation would interrupt the ongoing gesture-speech integration within a certain time window of the target brain location, providing direct evidence for the involvement of the TW in the gesture-speech priming effect, thus offering straightforward vision over the multisensory gesture-speech integration.

**Figure 1.**
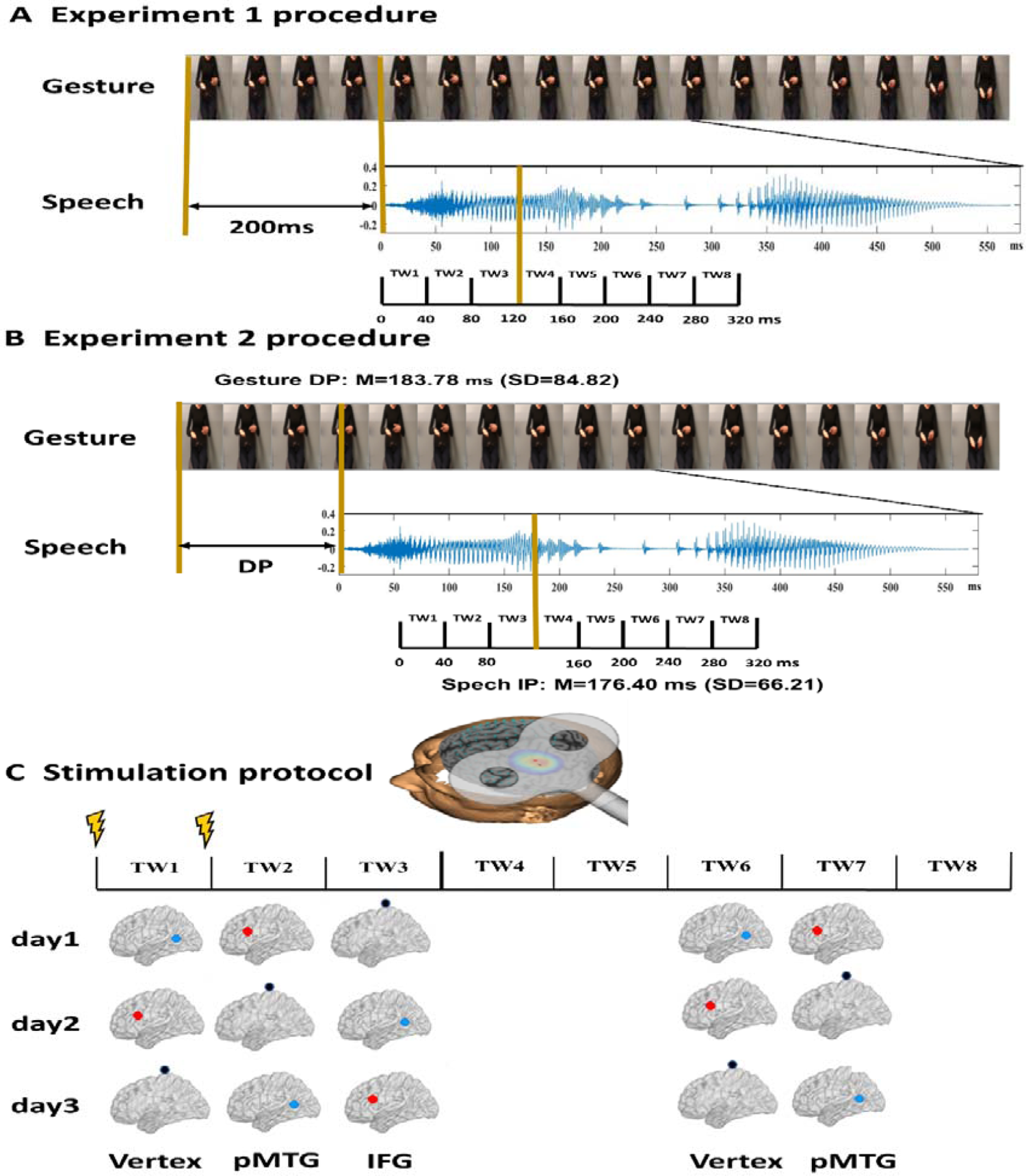
Experimental design and stimulus characteristics. (**A**) Procedure of Experiment 1. Twenty gestures were paired with 20 relevant speech stimuli. The onset of speech was set at 200 ms after the gesture. Eight time windows (TWs, duration = 40 ms) were segmented from the onset of speech. **(B)** Procedure of Experiment 2. Two gating studies were executed to define the minimal length of each gesture and speech required for semantic identification, namely, the DP of gesture (mean =183.78 ms, SD = 84.82) and the IP of speech (mean =76.40 ms (SD = 66.21). The onset of speech was set at the gesture DP. Eight TWs were segmented relative to the speech IP. (**C**) TMS protocol. Among the eight TWs, five (TW1, TW2, TW3, TW6, and TW7) were chosen based on the significant results of our prior study (Zhao et al., 2021). Double-pulse TMS was delivered over each of the TWs of either the pMTG, the IFG, or the vertex in a Latin-square balanced order.

## Materials and methods

### Apparatus and stimuli

Twenty gestures (length = 1771.00 ms, SD = 307.98) with 20 semantically congruent speech signals (length = 447.08 ms, SD = 93.48) were used in the present study. The stimuli set was recorded from two native Chinese speakers (1 male, 1 female) and validated with 30 participants by replicating the semantic congruency effect (reaction time (RT) difference between gesture-speech semantically congruent and semantically incongruent conditions), an index of the degree of gesture-speech integration. The results showed a significantly larger reaction time when participants were asked to judge the gender of the speaker if gestures contained incongruent semantic information with speech (a ‘cut’ gesture paired with speech word ‘喷pen1 (spray)’: mean = 554.51 ms, SE = 11.65) relative to when they were semantically congruent (a ‘cut’ gesture paired with ‘剪jian3 (cut)’ word: mean = 533.90 ms, SE = 12.02). (For details, see (Zhao et al., 2021).

Additionally, two separate pretests with 30 subjects in each (pretest 1: 16 females, aged 18-30 years, SD = 10.16; pretest 2: 15 females, aged 18-27 years, SD=8.94) were conducted to determine the minimal length of each gesture and speech required for semantic identification, namely, the DP of the gesture and the IP of speech. Using the gating paradigm (Obermeier and Gunter, 2015), participants were presented with segments of gesture and speech of increasing duration of 40 ms and were asked to infer what was described with a single action word. On average, the DP of gestures (mean =183.78 ms, SD =.84.82) and that of speech (Mean =176.40 ms, SD =66.21) were calculated (for details, see (Zhao et al., 2021).

To specify the priming stage of gesture upon speech, double-pulse TMS at an intensity of 50% of maximum stimulator output was delivered ‘online’ over either the left IFG or the left pMTG. Five 40-ms TWs that were selectively disrupted during gesture-speech integration were selected (Zhao et al., 2021). In Experiment 1, speech onset was set at 200 ms after the gesture stroke, and TWs were divided from the onset of speech (Zhao et al., 2018), i.e., TW1: 0∼40 ms, TW2: 40∼80 ms, TW3: 80∼120 ms, TW6: 200∼240 ms, TW7: 240∼280 ms of speech onset (**Figure 1A**). In Experiment 2, speech onset occurred at the DP of the gesture, and the five TWs were sorted relative to the speech IP (TW1: −120 ∼ −80 ms, TW2: −80 ∼ −40 ms, TW3: −40∼0 ms, TW6: 80∼120 ms, TW7: 120∼160 ms of speech IP) (Zhao et al., 2021) (**Figure 1B**).

To eliminate the effect caused by stimuli, each of the 160 gesture-speech pairs underwent TMS in each of the 5 TWs, leading to 800 trials in total. The 800 trials were further split into 10 blocks. Participants completed the 10 blocks on 3 different days that were 5-7 days apart to avoid fatigue. In a 3, 3, 4 order, one region was stimulated by double-pulse TMS in each block. The order of blocks and region being stimulated were balanced using a Latin square design (**Figure 1C**).

During the variable intertrial interval (ITI) of 0.5 to 1.5 sec, the stimuli were presented using Presentation software (Version 17.2, www.neurobs.com) in a pseudorandom order. The experimenter explained to participants that they would be presented with a number of videos with double-pulse TMS stimulation on. They were informed that the gender of the person they saw on the screen and the gender of the voice they heard might be different or might be the same. They were asked to indicate the gender of the voice they heard by pressing one of two buttons on the keyboard (key assignment was counterbalanced across participants). Participants’ reaction time, which was recorded relative to the onset of speech and the button being pressed, was recorded. Participants were allowed to have a break after each block and were told to press any button if they were ready to continue. Before the experiment began, participants were asked to perform 16 training trials to become accustomed to the experimental procedure.

We focused our analysis on the effect of semantic congruency and its interactions with the time window factor and the TMS effect factor (active-TMS minus Vertex-TMS) to determine at which TW the magnitude of the semantic congruency effect was reduced when activity in the pMTG or pMTG was stimulated relative to vertex stimulation. We also used the effect of gender congruency (the RT difference between gender incongruent and gender congruent conditions) as a control effect, with the assumption that double-pulse TMS stimulation would have an effect only on semantic congruency.

Accordingly, repeated measures analysis of variance (ANOVA) over each of the TMS effects (pMTG-Vertex and IFG-Vertex) on the RTs was conducted separately with respect to the effect of five TWs and its interaction with either semantic congruency (semantic congruent vs. semantic incongruent) or gender congruency (gender congruent vs. gender incongruent). Additionally, one-sample t tests with false discovery rate (FDR) correction of the TMS effect over the five TWs on either the semantic congruency effect or the gender congruency effect were implemented.

### TMS protocol

A Magstim Rapid² stimulator (Magstim, UK) was used to deliver the double-pulse TMS via a 70 mm figure-eight coil. The stimulation sites of the left IFG (−62, 16, 22) and the left pMTG (−50, −56, 10) corresponding to Montreal Neurological Institute (MNI) coordinates were identified in a quantitative meta-analysis of fMRI studies on iconic gesture-speech integration (for details, see (Zhao et al., 2018). The vertex was used as a control site.

To enable image-guided TMS navigation, high-resolution (1 × 1 × 0.6 mm) T1-weighted anatomical MRI scans of each participant were acquired at the Beijing MRI Centre for Brain Research using a Siemens 3T Trio/Tim Scanner. Frameless stereotaxic procedures (BrainSight 2; Rogue Research Inc, Montreal, Canada) were used for online checking of stimulation during navigation. To ensure precise stimulation of each target region for each participant, individual anatomical images were manually registered by identifying the anterior and posterior commissures. Subject-specific target regions were defined by trajectory markers using the MNI coordinate system. The angles of the markers were checked and adjusted to be orthogonal to the skull during neuronavigation.

## Experiment 1

### Method

#### Participants

Eighty participants (10 females, age 21-36, SD = 4.4) participated in Experiment 1. All were native Chinese speakers, were right-handed according to the Edinburgh Handedness form (laterality quotient (LQ) = 83.68, SD = 18.33), had normal or corrected-to-normal vision and were screened for TMS suitability using a medical questionnaire. Before the experiment, all signed consent forms approved by the Ethics Committee of the Department of Psychology, Chinese Academy of Sciences and were allowed for rTMS stimulation according to the TMS Subject Questionnaire. Participants were paid¥100 per hour for their participation.

## Results

All incorrect responses (518 out of the total number of 14400, 3.60% of trials) were excluded. To eliminate the influence of outliers, a 2SD trimmed mean for every participant in each session was conducted. Overall, larger RTs were reported for semantically incongruent trials (mean = 532, SE = 19.12) than for congruent trials (mean = 514, SE = 18.01). Furthermore, there were longer reaction times when speech and gestures were produced by different genders (mean = 536, SE = 18.72) than when they were produced by the same gender (mean = 509, SE = 18.60).

However, there was no significant interaction of stimulation site by semantic congruency (*F* _(1.751, 29.760)_ = .094, *p* = .887, ηp^2^ = .005), nor a significant interaction of stimulation site by TW by semantic congruency (*F* _(5.521, 93.856)_ = .469, *p* = .816, ηp^2^ = .027), as obtained from a 3 (stimulation site) × 5 (TW) × 2 (semantic congruency) × 2 (gender congruency) repeated-measures ANOVA. There was also no modulation of the gender congruency effect, as shown by the ANOVA, revealing no significant interactions, either for the interaction of stimulation site by gender Congruency (*F*_(1.995, 33.914)_ = 2.317, *p* = .114, ηp^2^ = .120), the interaction of stimulation site by TW by gender Congruency (*F*_(5.209, 88.545)_ = .307, *p* = .914, ηp^2^ = .018), or the four-way interaction of stimulation site by TW by semantic congruency by gender congruency (*F* _(4.875, 82.870)_ = .726, *p* = .603, ηp^2^ = .041). The results suggested that double-pulse TMS stimulation over a specific brain region did not modulate the magnitude of the semantic congruency effect or the gender congruency effect across the five time windows.

A 5 (TW) × 2 (semantic congruency) repeated-measures ANOVA on the TMS effect over the pMTG (pMTG - Vertex) revealed no significant main effect of semantic congruency (*F* _(1, 17)_ = 1.954, *p* = .180, η_p_^2^ = .103). There was no significant main effect of TW (*F*_(2.924, 49.709)_ = .593, *p* = .618, η_p_^2^ = .034) or a significant interaction of semantic congruency with TW (*F*_(2.862, 48.658)_ = .176, *p* = .950, η_p_^2^ = .010) (**Figure 2A**). Similar results were reported when the activity of the IFG was interrupted by double-pulse TMS, as shown by the nonsignificant main effect of semantic congruency (*F*_(1, 17)_ = 1.175, *p* = .293, η_p_^2^ = .065) and TW (*F*_(3.592, 61.067)_ = .505, *p* = .732, η_p_^2^ = .029) and the nonsignificant interaction of semantic congruency with TW (*F*_(3.178, 54.031)_ = .116, *p* = .976, η_p_^2^ = .007) (**Figure 2B**). Taken together, these results indicated that when activity of various time periods of IFG or pMTG was disrupted relative to the vertex, the gesture-speech semantically congruent trials and the semantically incongruent trials were not modulated differently.

**Figure 2.**
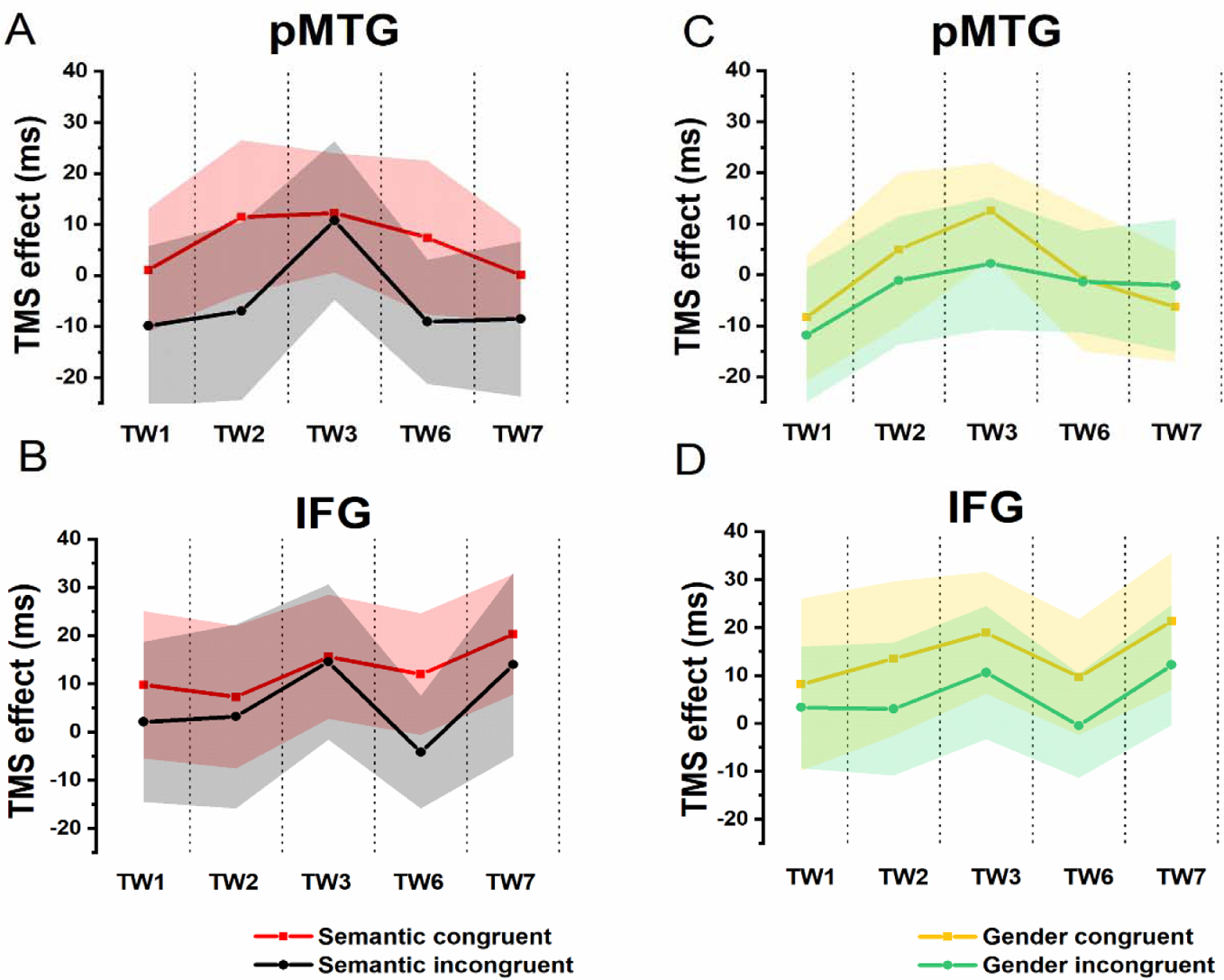
TMS effects on congruent and incongruent conditions in the semantic and gender domains. (A and B) RTs of TMS effects (active-TMS minus vertex-TMS) on the semantically congruent (red) and semantically incongruent (black) conditions when activity of the 5 TWs of the pMTG (A) and IFG (B) were interrupted. (C and D) TMS effect on RT of the gender congruent (yellow) and gender incongruent (green) conditions for stimulating the pMTG (C) and IFG (D) in the 5 TWs.

To directly indicate the TMS disruption over the semantic congruency effect (RT_semantically incongruent_ – RT_semantically congruent_), a one-sample t test with FDR correction was conducted over the five TWs. The results showed no significant TMS disruptions of the semantic congruency effect during the stimulation of either the pMTG (TW1: (*t*_(17)_ = .805, FDR-corrected *p* = .216, Cohen’s d = .190), TW2: (*t*_(17)_ = .963, FDR-corrected *p* = .174, Cohen’s d = .227), TW3: (*t*_(17)_ = .077, FDR-corrected *p* = .470, Cohen’s d = .018), TW6: (*t*_(17)_ = 1.041, FDR-corrected *p* = .156, Cohen’s d = .245), and TW7: (*t*_(17)_ = .671, FDR-corrected *p* = .256, Cohen’s d = .158)) or the IFG (TW1: (*t*_(17)_ = .572, FDR-corrected *p* = .288, Cohen’s d = .135), TW2: (*t*_(17)_ = .208, FDR-corrected *p* = .419, Cohen’s d = .049), TW3: (*t*_(17)_ = .069, FDR-corrected *p* = .473, Cohen’s d = .016), TW6: (*t*_(17)_ = 1.368, FDR-corrected *p* = .095, Cohen’s d = .322), and TW7: (*t*_(17)_ = .315, FDR-corrected *p* = .378, Cohen’s d = .074)) (**Figure 3A**).

**Figure 3.**
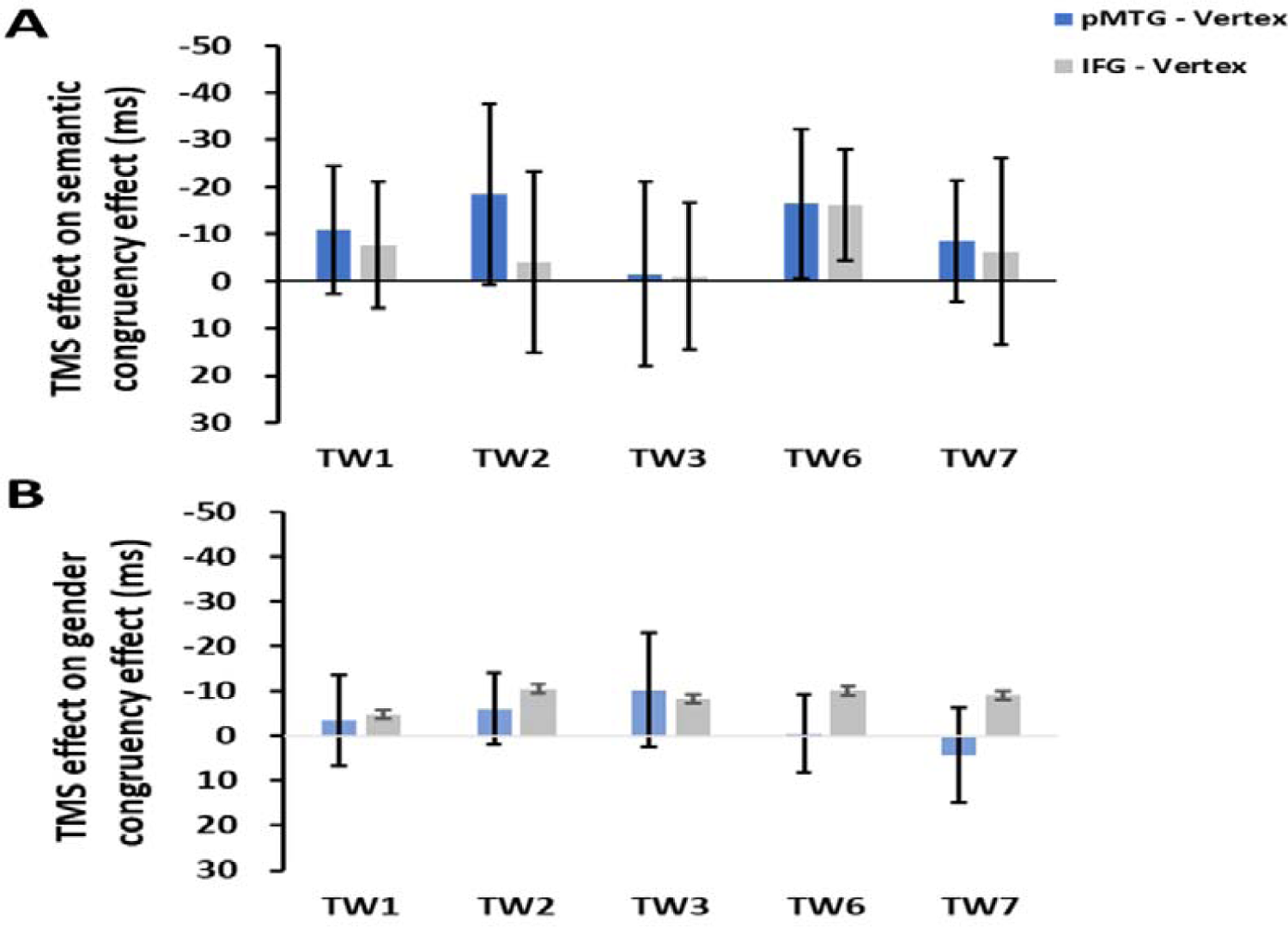
TMS effects of the pMTG (pMTG-TMS minus vertex-TMS, blue) and IFG (IFG-TMS minus vertex-TMS, gray) on the semantic congruency effect (RT difference between semantically incongruent and semantically congruent pairs, A) and the gender congruency effect (RT difference between gender incongruent and gender congruent pairs, B).

There was no modulation of the gender congruency effect, as shown by the 5 (TW) × 2 (gender congruency) repeated-measures ANOVA of the TMS effect of pMTG, revealing no significant main effects (gender congruency: (*F*_(1, 17)_ = .421, *p* = .525, η_p_^2^ = .024), TW: (*F*_(3.150, 53.543)_ = .606, *p* = .621, η_p_^2^ = .034)) nor a significant interaction (gender congruency * TW: (*F*_(3.185, 54.149)_ = .311, *p* = .829, η_p_^2^ = .018)) (**Figure 2C**). Another repeated-measures ANOVA of the TMS effect of IFG reported similar nonsignificant TMS modulation results: (gender congruency: (*F*_(1, 17)_ = 2.776, *p* = .114, η_p_^2^ = .140), TW: (*F*_(3.513, 59.720)_ = .395, *p* = .787, η_p_^2^ = .023), gender congruency * TW: (*F*_(3.509, 59.649)_ = .060, *p* = .993, η_p_^2^ = .004)) (**Figure 2D**). This result was followed by nonsignificant one-sample t test of TMS disruption over the gender congruency effect for both the pMTG (TW1: (*t*_(17)_ = .346, FDR-corrected *p* = .367, Cohen’s d = .082), TW2: (*t*_(17)_ = .769, FDR-corrected *p* = .226, Cohen’s d = .181), TW3: (*t*_(17)_ = .813, FDR-corrected *p* = .214, Cohen’s d = .192), TW6: (*t*_(17)_ = .059, FDR-corrected *p* = .477, Cohen’s d = .014, and TW7: (*t*_(17)_ = −.393, FDR-corrected *p* = .350, Cohen’s d = −.089) and the IFG (TW1: (*t*_(17)_ = .436, FDR-corrected *p* = .334, Cohen’s d = .103), TW2: (*t*_(17)_ = 1.108, FDR-corrected *p* = .142, Cohen’s d = .261), TW3: (*t*_(17)_ = .827, FDR-corrected *p* = .210, Cohen’s d = .195), TW6: (*t*_(17)_ = 1.229, FDR-corrected *p* = .118, Cohen’s d = .290, and TW7: (*t*_(17)_ = .885, FDR-corrected *p* = .194, Cohen’s d = .208) (**Figure 3B**).

## Experiment 2

### Method

#### Participants

Twenty-two participants (12 females, age 20-36, SD= 4.28) participated in Experiment 2. All of the participants were native Chinese speakers. Before the experiment, all signed consent forms were approved by the Ethics Committee of the Department of Psychology and were paid for their participation.

## Results

All incorrect responses (764 out of the total number of 17600, 4.34% of trials) were excluded. A 2SD trimmed mean for every participant in each session was conducted to eliminate the influence of outliers. The full pattern of results on the RTs revealed longer reaction times when speech and gestures were produced by different genders (mean = 508, SE = 12.91) than by the same gender (mean = 490, SE = 13.04). There were also larger RTs in semantically incongruent trials (mean = 507, SE = 12.96) than in congruent trials (mean = 491, SE = 12.98). The 3 (stimulation site) × 5 (TW) × 2 (semantic congruency) × 2 (gender congruency) repeated-measures ANOVA revealed a significant interaction of stimulation site by TW by semantic congruency (*F*_(5.177, 108.712)_ = 2.061, *p* = .042, ηp^2^ = .089). Meanwhile, there was no significant interaction of stimulation site by TW by gender congruency (*F*_(4.986, 104.702)_ = .450, *p* = .812, ηp^2^ = .021). Together, the results suggest that double-pulse TMS over different brain locations of various TWs would selectively impact only the semantic congruency effect, but not the gender congruency.

Additionally, the 5 (TW) × 2 (semantic congruency) repeated-measures ANOVA on the TMS effect over pMTG (pMTG - Vertex) revealed a significant interaction of TW with semantic congruency (*F*_(2.605, 54.701)_ = 3.951, *p* = .042, η_p_^2^ = .143). This illustrated that the magnitude of semantic congruency was modulated by different TWs when double-pulse TMS stimulation was applied over these TWs in different brain areas (pMTG or Vertex). Simple effect analysis showed a significant difference in terms of TMS disruption of the pMTG in the semantically incongruent condition compared to the semantically congruent condition in TW1 (*F*_(1, 21)_ = 5.414, *p* = .030, η_p_^2^ = .205), TW2 (*F*_(1, 21)_ = 9.547, *p* = .006, η_p_^2^ = .313), TW6 (*F*_(1, 21)_ = 14.799, *p* <.001, η_p_^2^ = .413) and TW7 (*F*_(1, 21)_ = 8.111, *p* = .010, η_p_^2^ = .279) (**Figure 4A**).

**Figure 4.**
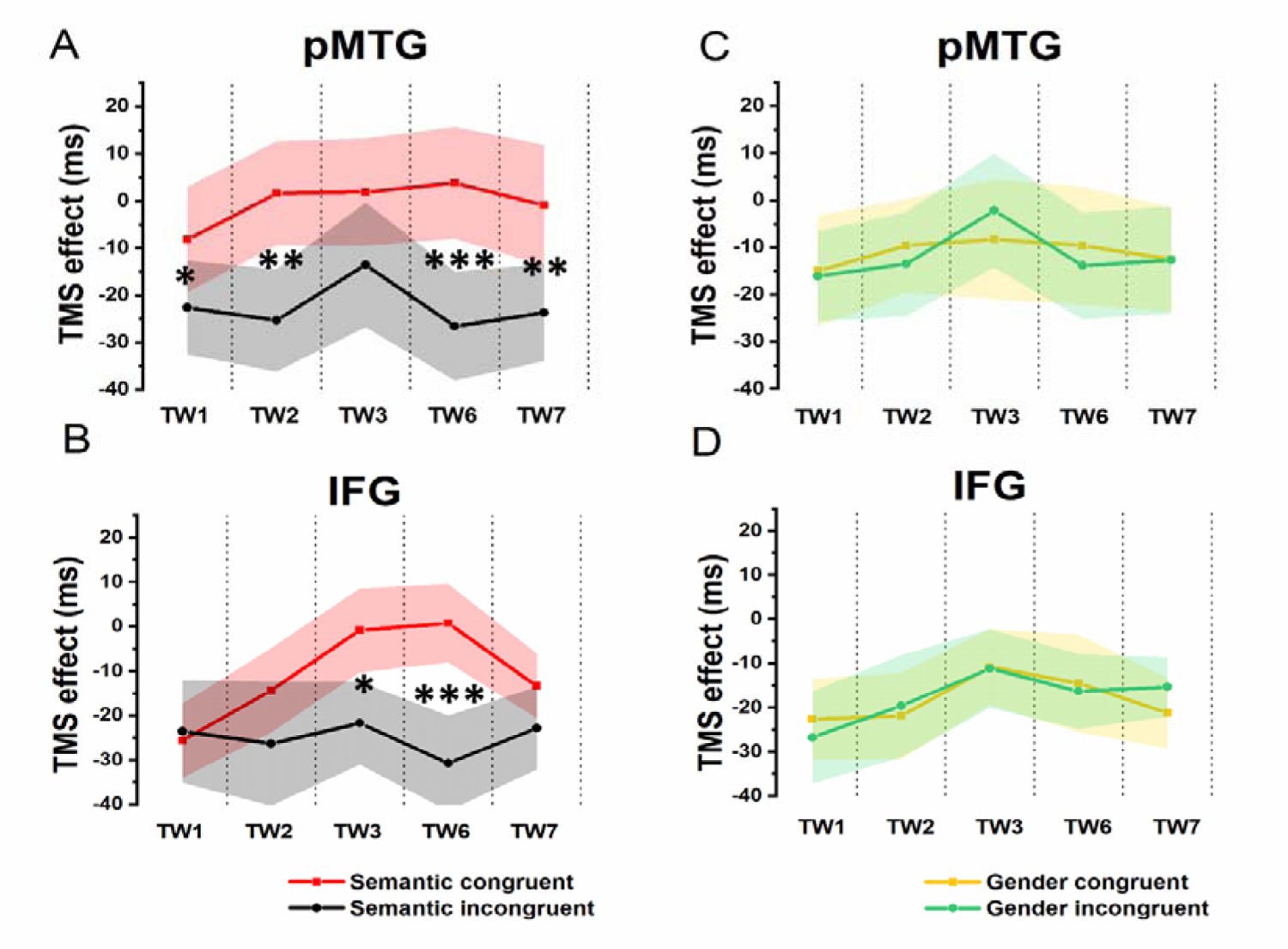
TMS effects on congruent and incongruent conditions in the semantic and gender domains. (A and B) RTs of TMS effects (active-TMS minus vertex-TMS) on the semantically congruent (red) and semantically incongruent (black) conditions when activity of the 5 TWs of the pMTG (A) and IFG (B) were interrupted. (C and D) TMS effect on RT of the gender congruent (yellow) and gender incongruent (green) conditions for stimulating the pMTG (C) and IFG (D) in the 5 TWs.

A significant interaction of TW with semantic congruency (*F*_(3.171, 66.601)_ = 2.749, *p* = .047, η_p_^2^ = .116) was also reported in the 5 (TW) × 2 (semantic congruency) repeated-measures ANOVA on the TMS effect over the IFG (IFG - Vertex). Simple effect analysis showed a significant semantic difference in terms of the TMS effect over the IFG in TW3 (*F*_(1, 21)_ = 5.337, *p* = .031, η_p_^2^ = .203) and TW6 (*F*_(1, 21)_ = 20.089, *p* <.001, η_p_^2^ = .489) (**Figure 4B**).

The direct site- and TW-specific TMS effect on the semantic congruency effect of one-sample t tests showed significant TMS disruptions during the stimulation of the pMTG in TW1 (*t*(21) = 2.327, FDR-corrected *p* = .015, Cohen’s d = .496), TW2 (*t*(21) = 3.019, FDR-corrected *p* = .003, Cohen’s d = .659), TW6 (*t*(21) = 3.847, FDR-corrected *p* <.001, Cohen’s d = .820) and TW7 (*t*(21) = 2.848, FDR-corrected *p* = .005, Cohen’s d = .607). Similar TMS impairment of the semantic congruency effect was found during the stimulation of the IFG in TW3 (*t*(21) = 2.310, FDR-corrected *p* = .016, Cohen’s d = .493) and TW6 (*t*(21) = 4.482, FDR-corrected *p* <.001, Cohen’s d = .956) (**Figure 5A**).

**Figure 5.**
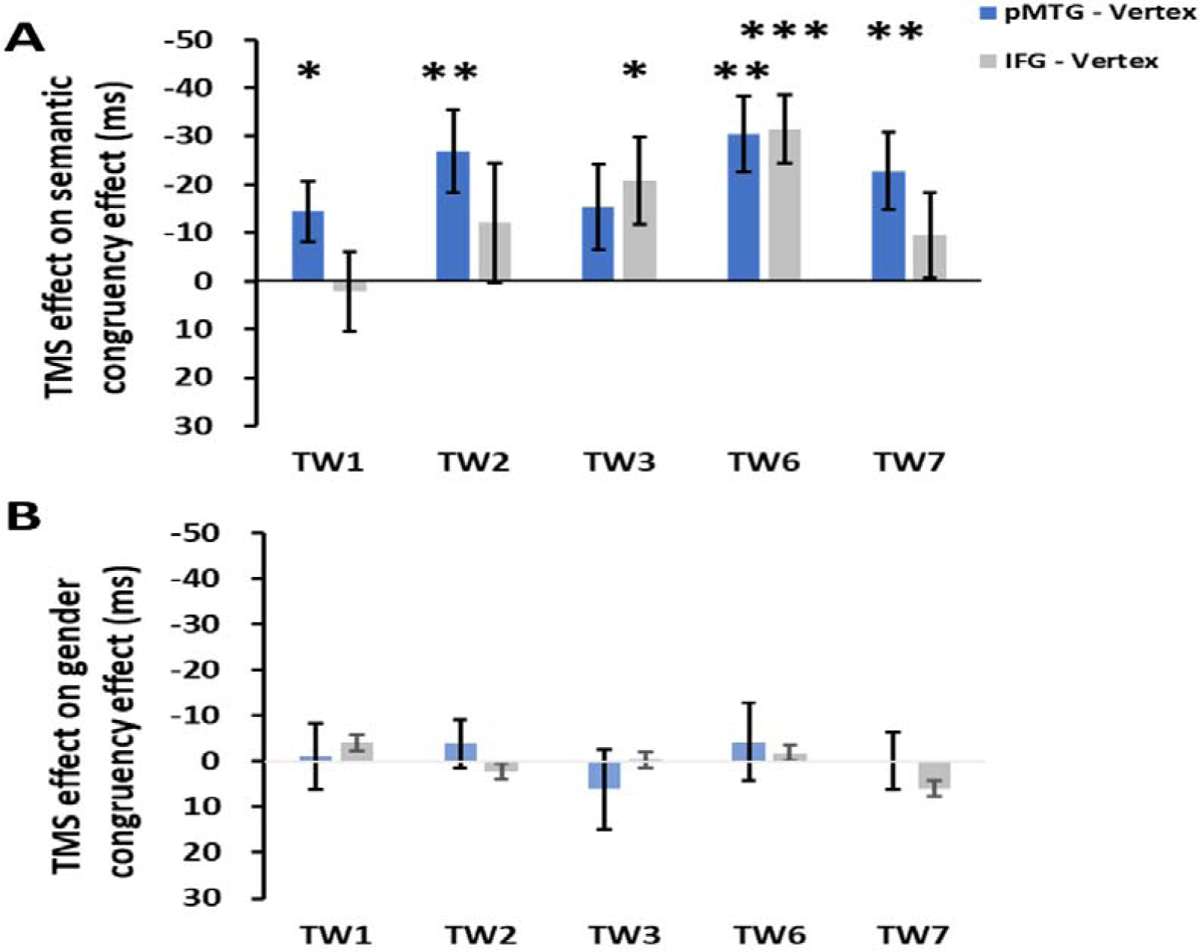
TMS effects over the pMTG (pMTG-TMS minus vertex-TMS, blue) and IFG (IFG-TMS minus vertex-TMS, gray) on the semantic congruency effect (RT difference between semantically incongruent and semantically congruent pairs, A) and the gender congruency effect (RT difference between gender incongruent and gender congruent pairs, B).

There was no modulation of the gender congruency effect, as the ANOVA of the TMS effect over both the pMTG and IFG failed to show any other significant effects. Regarding the TMS effect over the pMTG, the following results were found: main effect of gender congruency (*F*_(1, 21)_ = .076, *p* = .786, η_p_^2^ = .004), main effect of TW (*F*_(3.111, 65.337)_ = .911, *p* = .443, η_p_^2^ = .042), and interaction of TW with gender congruency (*F*_(3.062, 64.292)_ = .292, *p* = .835, η_p_^2^ = .014) (**Figure 4C**). Regarding the TMS effect over the IFG, the following results were found: main effect of gender congruency (*F*_(1, 21)_ = .012, *p* = .915, η_p_^2^ = .001), main effect of TW (*F*_(3.399, 71.379)_ = 2.119, *p* = .098, η_p_^2^ = .092), and interaction of TW with gender congruency (*F*_(3.358, 70.516)_ = .420, *p* = .761, η_p_^2^ = .020) (**Figure 4D**). There was no significant TMS effect reported by the one-sample t test for either the pMTG (TW1: (*t*_(21)_ = .153, FDR-corrected *p* = .440, Cohen’s d = .033), TW2: (*t*_(21)_ = .734, FDR-corrected *p* = .235, Cohen’s d = .157), TW3: (*t*_(21)_ = .705, FDR-corrected *p* = .244, Cohen’s d = .150), TW6: (*t*_(21)_ = .501, FDR-corrected *p* = .311, Cohen’s d = .107), and TW7: (*t*_(21)_ = .021, FDR-corrected *p* = .492, Cohen’s d = .005)) or the IFG (TW1: (*t*_(21)_ = .596, FDR-corrected *p* = .279, Cohen’s d = .127), TW2: (*t*_(21)_ = -.366, FDR-corrected *p* = .359, Cohen’s d = -.078), TW3: (*t*_(21)_ = .057, FDR-corrected *p* = .478, Cohen’s d = .02), TW6: (*t*_(21)_ = .258, FDR-corrected *p* = .400, Cohen’s d = .055), and TW7: (*t*_(21)_ = −1.267, FDR-corrected *p* = .110, Cohen’s d = -.270)) (**Figure 5B**).

## Discussion

By applying double-pulse TMS stimulation over the integration node of the pMTG and IFG during various TWs of gesture-speech integration, the present study directly tested how gestures prime speech. Crucially, the present study observed a significantly interrupted semantic congruency effect only when gestures were primed semantically with speech and when the TWs were segmented in accordance with comprehensive speech stages (Experiment 2). When there was a fixed time advance of gesture over speech, and TWs were split from the onset of speech, as reported in Experiment 1, there was no significant disruption of gesture-speech integration. Our findings provide the first causal evidence of how the automatic priming effect of gestures on speech takes place and thus provide a glimpse into the integration of multisensory gesture-speech semantic information.

Using the gesture-speech semantic congruency effect as an index of the degree of integration (Kelly et al., 2010a; Zhao et al., 2018), the present study provided direct evidence for the lexical priming effect of gestures on speech. We hypothesized that this priming effect is how gestures and speech ‘integrate’ with each other. Additionally, our results showed that there was no semantic congruency effect in Experiment 1, yet this effect became significant when we shifted the TWs in accordance with the speech IP to clearly identify the processing stage of gestures and speech in Experiment 2. We conceived that gestures need to be comprehended to have a priming effect on speech, in which the information conveyed in gestures is encoded as a whole (McNeill, 1992) and based on the goal rather than the individual movement of the action form (Umilta et al., 2008; Rizzolatti et al., 2009). In other words, the representation of gestures should be established before these gestures can act as an equivalent lexical formation to further influence lexical selection.

Two stages were differentiated for the priming effect of gestures upon speech: before speech reaches its semantic IP and after speech has been clearly understood. Here, we defined two stages of gesture-speech interaction: semantic priming of gestures onto the phonological processing of speech and unification of gesture and speech semantics to form context-appropriate semantic representations. The two processing stages in gesture-speech integration proposed in the present study are consistent with previous findings (Drijvers et al., 2018; He et al., 2018). In a study by (Kelly et al., 2004), when information contained in the preceding gesture and the following speech word was incongruent, there were early N1-P1 and P2 sensory effects followed by an N400 semantic effect; this was interpreted to indicate that gesture influenced word comprehension first at the level of sensory/phonological processing and later at the level of semantic processing. Of particular note, the finding that gestures prime speech first in a sensory processing stage, followed by a lexico-semantic processing stage, replicates our previous study (Zhao et al., 2021) using the same TMS protocol. As a step forward, by contrasting different segmentations of the gesture-speech integration period in reference to either speech onset or speech comprehension processes in two separate experiments, the present study offered straightforward proof for the two stages during integration, thus settling the debate on how gestures prime speech.

The theoretical model of gesture-speech integration has been investigated extensively (Bernardis and Gentilucci, 2006; Ozyurek et al., 2007; Kelly et al., 2010b). In particular, (Krauss, 2000) proposed a lexical gesture process model to explain the facilitation effect of gestures on speech. According to this model, information stored in memory is encoded in multiple ways, including both the visuospatial format of information arising from gestures and the conceptual format of speech information. The access of one format of information (gesture) will activate its representation that is stored in the visuo-spatial working memory (Wu and Coulson, 2014; Wu et al., 2022), and the activated representation (of gesture) will result in the spreading of activation of the related representations (speech). Using double-pulse TMS with high temporal resolution, the present study causally confirmed that activated gesture representation influences speech lexical selection. During the process, the motoric feature of the gesture is selected by the kinesthetic monitor, and then the selected information is transferred into the phonological encoder of speech, where it facilitates the lexical retrieval of words (Krauss, 2000). Next, the retrieved lexical information of speech is sustained and unified with the previously processed gesture to form a coherent meaning.

We further tested the sequential involvement of the pMTG and IFG in the two stages of gesture-speech integration, as reported by a feedforward connection from the pMTG to the IFG before the speech IP (TW1-TW3) and a feedback connection from the IFG to the pMTG after speech identification (TW6 and TW7). Consistent with a previous study (Zhao et al., 2021), the connection from the left pMTG to the left IFG in the prelexical stage was understood to enable semantic selection and activation, and the pathway from the left IFG to the left pMTG in the postlexical stage was explained as allowing the unification of multimodal information.

Notably, the experimental paradigm used in the present study provided an automatic paradigm to examine the relationship between gestures and speech. Our results showed that even if participants were not asked to pay attention to the information conveyed in gestures, the representation of gestures that had been learned by participants still affected the lexical retrieval of speech that cooccurred. Therefore, the hypothesis of the two-stage model proposed applies only when the effect between gesture and speech takes place in an automatic, obligatory way. Since gesture-speech interaction can also be modulated by some controlled processes (Holle and Gunter, 2007; Kelly et al., 2007), whether and how the two-stage model proposed here works need to be clarified in future studies.

In summary, by segmenting the gesture-speech integration process in reference to either time units or the semantic processing stage and applying double-pulse TMS upon each scale separately, the present study clarified the priming effect of gestures upon speech. For the first time, we reported that gesture-speech integration takes place after gestures have been clearly comprehended such that gestures first prime lexical retrieval during the speech sensory processing stage and then unify with speech in the lexico-semantic stage. This finding provides new insights into multimodal gesture-speech semantic integration.

## Notes

### Competing Interest Statement

The authors have declared no competing interest.

